# Genome-wide discovery of SLE genetic risk variant allelic enhancer activity

**DOI:** 10.1101/2020.01.20.906701

**Authors:** Xiaoming Lu, Xiaoting Chen, Carmy Forney, Omer Donmez, Daniel Miller, Sreeja Parameswaran, Ted Hong, Yongbo Huang, Mario Pujato, Tareian Cazares, Emily R. Miraldi, John P. Ray, Carl G. de Boer, John B. Harley, Matthew T. Weirauch, Leah C. Kottyan

**Affiliations:** Center for Autoimmune Genomics and Etiology, Cincinnati Children’s Hospital Medical Center, Cincinnati, Ohio, USA, 45229; Department of Pharmacology & Systems Physiology, University of Cincinnati, College of Medicine, Cincinnati, Ohio, USA, 45229; Division of Biomedical Informatics, Cincinnati Children’s Hospital Medical Center, Cincinnati, Ohio, USA, 45229; Division of Immunobiology, Cincinnati Children’s Hospital Medical Center, Cincinnati, Ohio, USA, 45229; Department of Pediatrics, University of Cincinnati, College of Medicine, Cincinnati, Ohio, USA, 45229; Broad Institute of Massachusetts Institute of Technology (MIT) and Harvard University, Cambridge, Massachusetts, USA, 02142; US Department of Veterans Affairs Medical Center, Cincinnati, Ohio, USA 45229; Division of Developmental Biology, Cincinnati Children’s Hospital Medical Center, Cincinnati, Ohio, USA, 45229; Division of Allergy and Immunology, Cincinnati Children’s Hospital Medical Center, Cincinnati, Ohio, USA, 45229

## Abstract

Genome-wide association studies of Systemic Lupus Erythematosus (SLE) nominate 3,073 genetic variants at 91 risk loci. To systematically screen these variants for allelic transcriptional enhancer activity, we constructed a massively parallel reporter assay (MPRA) library comprising 12,396 DNA oligonucleotides containing the genomic context around every allele of each SLE variant. Transfection into the Epstein-Barr virus-transformed B cell line GM12878 revealed 482 variants with enhancer activity, with 51 variants showing genotype-dependent (allelic) enhancer activity at 27 risk loci. Comparison of MPRA results in GM12878 and Jurkat T cell lines highlights shared and unique allelic transcriptional regulatory mechanisms at SLE risk loci. In-depth analysis of allelic transcription factor (TF) binding at and around allelic variants identifies one class of TFs whose DNA-binding motif tends to be directly altered by the risk variant and a second, larger class of TFs that bind allelically without direct alteration of their motif by the variant. Collectively, our approach provides a blueprint for the discovery of allelic gene regulation at risk loci for any disease and offers insight into the transcriptional regulatory mechanisms underlying SLE.

---

Systemic Lupus Erythematosus (SLE) is an autoimmune disease that can affect multiple organs, leading to debilitating inflammation and mortality^1^. Up to 150 cases are found per 100,000 individuals, and the limited treatment options contribute to considerable economic and social burden^1,2^. Epidemiological studies have established a role for both genetic and environmental factors in the development of SLE^2^. SLE has a relatively high heritability^3^. The vast majority of patients do not have a single disease-causing mutation (such as mutations in complement protein 1q); instead, genetic risk is accumulated additively through many genetic risk loci with modest effect sizes^4^.

Genome-wide association studies (GWASs) have identified 91 genetic risk loci that increase disease risk of SLE in a largely additive fashion^4^. Each SLE risk locus is a segment of the genome containing a polymorphic “tag” variant (i.e. the variant with the most significant GWAS p-value) and the genetic variants in linkage disequilibrium with the tag variant. The majority (68%) of the established SLE risk loci do not contain a disease associated coding variant that changes amino acid usage^5^. Instead, variants at these loci are found in non-coding regions of the genome such as introns, promoters, enhancers, and other intergenic areas. Enrichment of these variants in enhancers and at transcription factor (TF) binding sites^6,7^ implies that transcriptional perturbation may be a key to the development of SLE^8^. However, given the large number of candidate variants identified by GWASs, identification of the particular causal variant(s) remains challenging.

SLE is a complex disease that involves multiple cell types^2^. Previous systematic studies demonstrate that SLE risk loci are enriched for B cell specific genes^9^ and regulatory regions^10^. Established biological mechanisms further highlight a key role for B cells in SLE - as the autoantibody-secreting cell type, B cells are critical to the pathoetiology of SLE, a disease characterized by autoantibody production^11^. B cells also present self-antigens to T cells in the development of an autoantigen-focused (i.e. “self”) inflammatory response^12^. Meanwhile, Epstein-Barr virus (EBV)-infected B cells have been implicated in SLE, with patients having a greater number of EBV-infected B cells and a higher viral load than people without SLE^13,14^. In addition, EBV infection is significantly more prevalent in SLE cases than controls^15,16^, and EBV-encoded EBNA2 interactions with the human genome are concentrated at SLE risk loci in EBV-transformed B cell lines^10^. *In vitro*, EBV infection can transform B cells into a lymphoblastoid cell line (LCL)^17^. We have recently shown that histone mark and human and viral protein chromatin immunoprecipitation followed by sequencing (ChIP-seq) datasets from EBV-transformed B cell lines are highly and specifically enriched at SLE risk loci relative to non-EBV transformed B cell lines^9,10^. Given the above evidence, we chose the EBV-transformed B cell line GM12878 to study the effects of SLE risk variants at many SLE risk loci.

To systematically identify the SLE genetic risk variants that contribute to transcriptional dysregulation in the EBV-transformed B cell line GM12878, we designed and applied a massively parallel reporter assay (MPRA)^18-27^ (**Figure 1, Supplemental Figure 1**). MPRA extends standard reporter assays, replacing low-throughput luciferase with high-throughput mRNA expression detection. In this study, we used MPRA to simultaneously screen the full set of genome-wide significant SLE-associated genetic variants for effects on gene regulation. Using this experimental approach, we nominate 51 putative causal variants that result in genotype-dependent (allelic) transcriptional regulation. Comparison of MPRA results between GM12878 and the Jurkat T cell line reveals shared and cell-type specific allelic behavior. Integration of these data with TF binding site predictions and functional genomics data reveals two distinct mechanisms whereby TFs bind risk variants in an allelic manner - directly impacted by a given variant (i.e. the variant directly alters the TF’s DNA-binding site) or indirectly impacted by the variant (i.e. the variant alters the DNA binding of the TF’s physical interaction partner or modulates chromatin accessibility). Collectively, these results provide an important resource for understanding SLE disease risk mechanisms and reveal an important role for groups of TFs in the mediation of allelic enhancer activity at plausibly causal SLE risk variants in EBV-transformed B cells.

**Figure 1.**
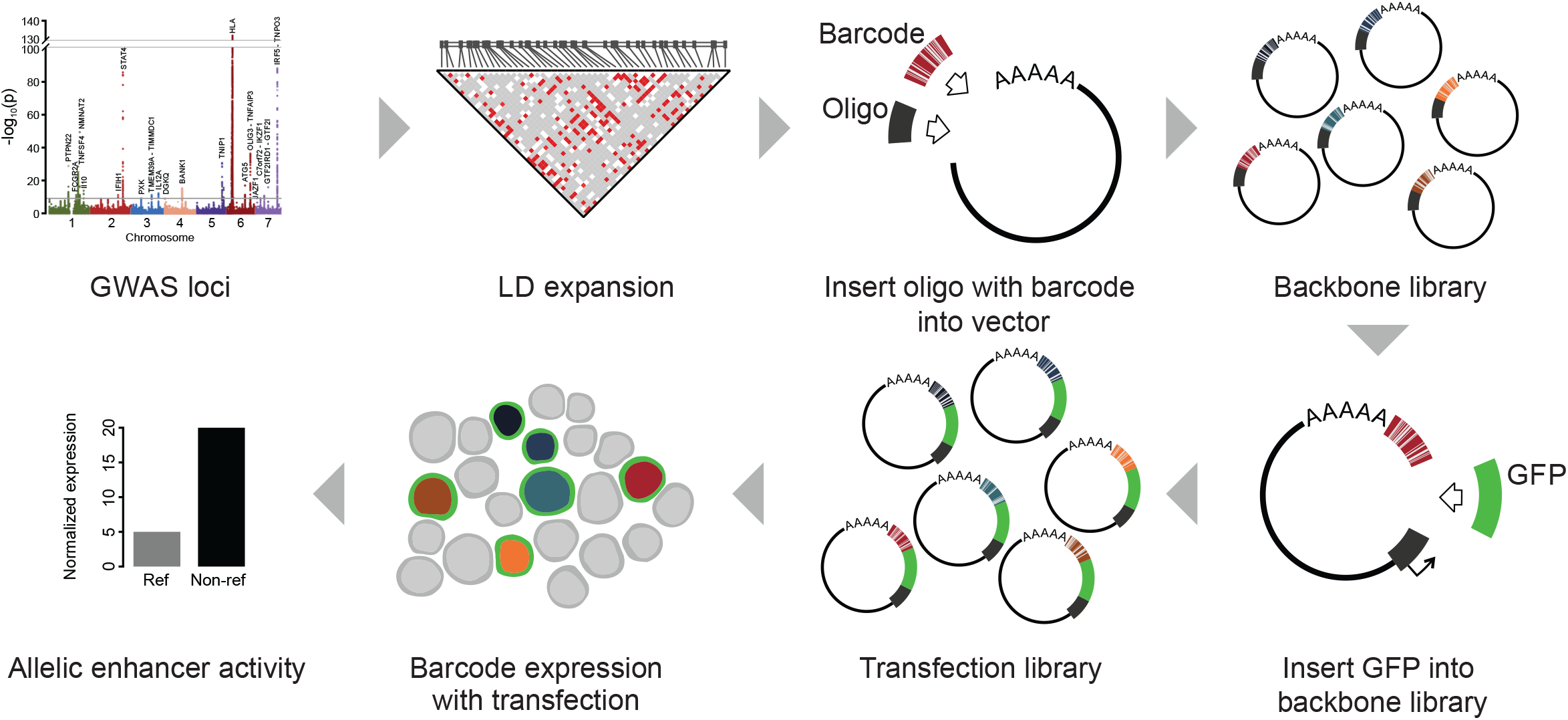
Massively Parallel Reporter Assays Workflow. See text for description.

## Results

### MPRA library design and quality control

We first collected all SLE-associated risk loci reaching genome-wide association significance (p<5×10^−8^) published through March 2018 **(Supplemental Data Set 1)**. Studies of all ancestral groups were included, and independent risk loci were defined as loci with lead (tag) variant at r^2^<0.2. For each of these 91 risk loci, we performed linkage disequilibrium (LD) expansion (r^2^>0.8) in each ancestry of the initial genetic association(s), to include all possible disease-relevant variants **(Supplemental Data Set 2)**. In total, this procedure identified 3,073 genetic variants. All established alleles of these variants were included, with 36 variants having three or more alleles. We also included 20 additional genetic variants from a previously published study^19^ as positive and negative controls to assess the library’s performance **(Supplemental Data Set 3)**.

For each variant, we generated a pair of 170 base pair (bp) DNA oligonucleotides (subsequently referred to as “oligos”) for each allele, with the variant located in the center and identical flanking genomic sequence across the alleles **(Supplemental Data Set 4)**. A total of 12,478 oligos (3,093 variants with 6,239 alleles) were synthesized. For barcoding, a pool of random 20mers were added to the oligo through PCR. Each unique barcode was matched with perfectly synthesized oligos. The number of unique barcodes per oligo had an approximately normal distribution with a median of 729 barcodes per oligo **(Supplemental Figure 2A, Supplemental Data Set 5)**. Only oligos with at least 30 unique barcodes were used for downstream analyses. A fragment containing a minimal promoter and an eGFP gene was inserted between the oligo and barcode to generate the MPRA transfection library. We note that the use of a minimal promoter allows us to effectively measure the ability of alleles to enhance, but not reduce, transcriptional activity. Three aliquots of the library were independently transfected into the EBV-transformed B cell line GM12878. We then used nucleic acid capture to enrich for eGFP mRNA and sequenced the barcode region. The normalized barcode ratio between the eGFP mRNA and the plasmid DNA was used to quantify the amount of enhancer activity driven by each oligo (see **Supplemental Note 1, Supplemental Figure 5F, G**). This mRNA to DNA ratio measures the enhancing effect of an allele on eGFP expression under the control of a minimal promoter **(Figure 1, Supplemental Figure 1)**. We observed strong correlation of enhancer activity between experimental replicates (mean pairwise Pearson correlation of 0.99) **(Supplemental Figure 2B, C, D)**. Likewise, calibration variants showed high accuracy, with 17 of the 20 variants matching the results of a previous study^19^ (87.5% sensitivity and 75% specificity), collectively demonstrating a robust experimental system **(Supplemental Data Set 3)**.

### Hundreds of SLE risk variants are located in genomic regions with enhancer activity in EBV-transformed B cells

Using the SLE MPRA library, we next identified genetic variants capable of driving enhancer activity in the EBV-transformed B cell line GM12878. An SLE risk variant was considered a candidate for enhancer activity if an oligo corresponding to any allele had significantly increased transcriptional regulatory activity compared to controls (see Methods). Not all statistically significant changes in transcriptional activity are necessarily biologically relevant – a highly consistent, but slight change in expression levels is statistically, but not biologically, meaningful. We therefore considered an oligo to have enhancer activity only when (1) the oligo had statistically significant enhancer activity (p_FDR_<0.05) and (2) we observed at least a 50% increase in transcriptional activity compared to the corresponding barcode counts in the plasmid control. Based on these criteria, 16% of SLE risk variants (482 variants, 853 alleles) demonstrated enhancer activity, henceforth referred to as “enhancer variants” (enVars) and “enhancer alleles” (enAlleles), respectively (**Figure 2A, Supplemental Data Set 6**).

**Figure 2.**
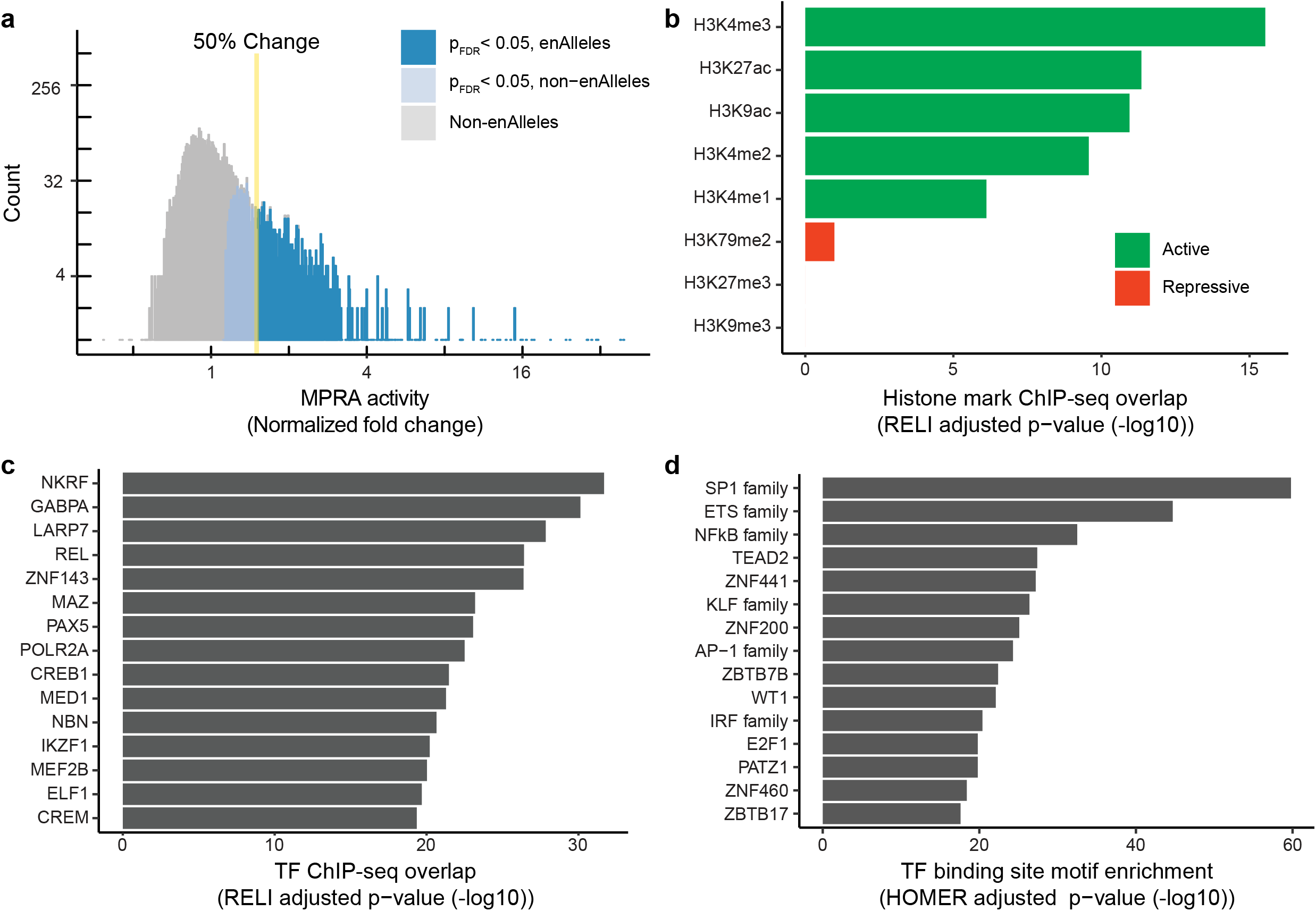
Regulatory activity of enhancer variants (enVars). a. Distribution of MPRA regulatory activity. The normalized fold-change of MPRA activity relative to plasmid control (X-axis) was calculated using DESeq2. Enhancer alleles (enAlleles) were identified as those alleles with significant activity relative to control (p_adj_ < 0.05) and at least a 50% increase in activity (see Methods). b. Enrichment of histone marks in GM12878 cell at enVars compared to non-enVars. p-values were estimated using RELI (see Methods). Full RELI results are provided in **Supplemental Data Set 9**. c. Enrichment of regulatory protein and transcription factor (TF) binding at enVars compared to non-enVars. The top 15 TFs (based on RELI p-values) that overlap at least 10% of enVars are shown. Full results are provided in **Supplemental Data Set 9**. d. TF binding site motif enrichment for enVars compared to non-enVars. p-values were obtained from HOMER using the full oligo sequences of enVars and non-enVars (see Methods). The top 15 enriched TF motif families are shown. Full results are provided in **Supplemental Data Set 10**.

We next explored the potential effects of enVars on gene expression. We connected each enVar to one or more genes using an approach that takes into account chromatin looping interactions, expression quantitative trait loci (eQTLs), and gene proximity (**Supplemental Data Set 2, Supplemental Data Set 7**) (see Methods). This approach identified 1,006 genes in total, which are enriched for expected SLE-related processes such as the interferon pathway, the antigen processing and presentation pathway, and cytokine-related pathways (**Supplemental Figure 3, Supplemental Data Set 8**), providing functional support for the enVars we identified.

Next, we searched for functional genomic features enriched within enVars relative to non-enVars using the RELI algorithm^10^. In brief, RELI estimates the significance of the intersection between an input set of genomic regions (e.g., enVars) and each member of a collection of functional genomics datasets (e.g., ChIP-seq for a particular histone mark or TF). For this analysis, we identified, curated, and systematically processed the 576 GM12878 ChIP-seq datasets available in the NCBI Gene Expression Omnibus (GEO) database (see Methods). Using RELI, we observed significant enrichment for overlap between enVars and multiple histone modification marks, including H3K4me3 (5.8-fold, p_corrected_ <10^−21^) and H3K27ac (2.0-fold, p_corrected_ <10^−13^) (**Figure 2B, Supplemental Data Set 9**). As expected, we did not identify enrichment for repressive marks such as H3K9me3 or H3K27me3^28^ (**Figure 2B, Supplemental Data Set 9**). Altogether, the genomic features present within enVars confirm that many SLE genetic risk loci likely alter transcriptional regulation in EBV-transformed B cells.

We next asked if the genomic binding sites of particular TFs were enriched within our enVars using RELI and the TF ChIP-seq datasets from GM12878. As expected, the enVars are highly enriched for ChIP-seq signal of TFs involved in regulation of the immune response, relative to variants lacking enhancer activity (**Figure 2C, Supplemental Data Set 9**). In particular, we found significant enrichment for all members of the NFκB TF family: REL/C-Rel (6.4-fold, p_corrected_ <10^−26^), NFKB1/p50 (3.0-fold, p_corrected_ <10^−18^), RELA/p65 (3.1-fold, p_corrected_ <10^−16^), RELB (2.7-fold, p_corrected_ <10^−10^), and NFKB2/p52 (2.2-fold, p_corrected_ <10^−7^). These results are consistent with our previous findings that altered binding of NFκB TFs is likely an important mechanism conferring SLE risk^10^. We also found significant enrichment for other TFs that have been previously implicated in SLE pathogenesis, such as PAX5^29^, MED1^30^, IKZF1^31^, ELF1^32^ and the EBV-encoded EBNA2 transactivator^10^ (**Figure 2C, Supplemental Data Set 9**). As a complementary approach, we next assessed enrichment for TF binding site motifs in the enAllele DNA sequences using HOMER^33^ and motifs contained in the Cis-BP database^34^ (see Methods). This analysis also revealed enrichment of multiple TF families with known roles in SLE, including ETS, NFκB, and IRF^3^ (**Figure 2D, Supplemental Data Set 10**). Many of these same TFs also have enriched ChIP-seq peaks at SLE risk loci^10^. Collectively, these results indicate that particular TFs tend to not only concentrate at SLE risk loci^10^, but also concentrate at alleles capable of driving gene expression in EBV-transformed B cells.

### MPRA identifies 51 SLE risk variants with allelic enhancer activity in EBV-transformed B cells

We next used our MPRA library to identify SLE genetic risk variants that drive allele-dependent (allelic) enhancer activity. Allelic activity was assessed for each enVar by comparing enhancer activity between each pairs of alleles. We considered a SLE variant allelic if (1) at least one of its alleles is an enAllele; (2) we observed significant genotype-dependent activity using Student’s *t*-test^19,35^ (see **Supplemental Note 2, Supplemental Figure 6**); and (3) the oligos had more than a 25% change between any pair of alleles. Using these criteria, we identified 51 SLE risk variants (11% of enVars, 1.7% of all SLE risk variants) as allelic enVars in GM12878 (**Figure 3A, Supplemental Data Set 11**). For 31 of these 51 allelic enVars, the risk allele decreased enhancer activity relative to the non-risk allele, which is statistically indistinguishable from the 20 variants with increased risk allele activity (p=0.1). Three of the allelic enVars can also alter the amino acid sequence of proteins – rs1059702 (IRAK1), rs1804182 (PLAT), and rs3027878 (HCFC1), consistent with previous studies identifying dual-use codons in the human genome^36^. Collectively, these 51 variants represent causal variant candidates for 27 SLE risk loci (30% of all tested loci) (**Supplemental Data Set 12**). For these 27 risk loci, our approach reduced the number of potential causal variants with allelic activity in GM12878 from an average of 84 variants to an average of two variants per risk locus (**Figure 3B**). For example, at 17q12 (marked by rs8079075), we reduce the candidate causal variant set from 249 to one, with the rs112569955 “G” risk allele showing a 36% increase in enhancer activity compared to the “A” non-risk allele.

**Figure 3.**
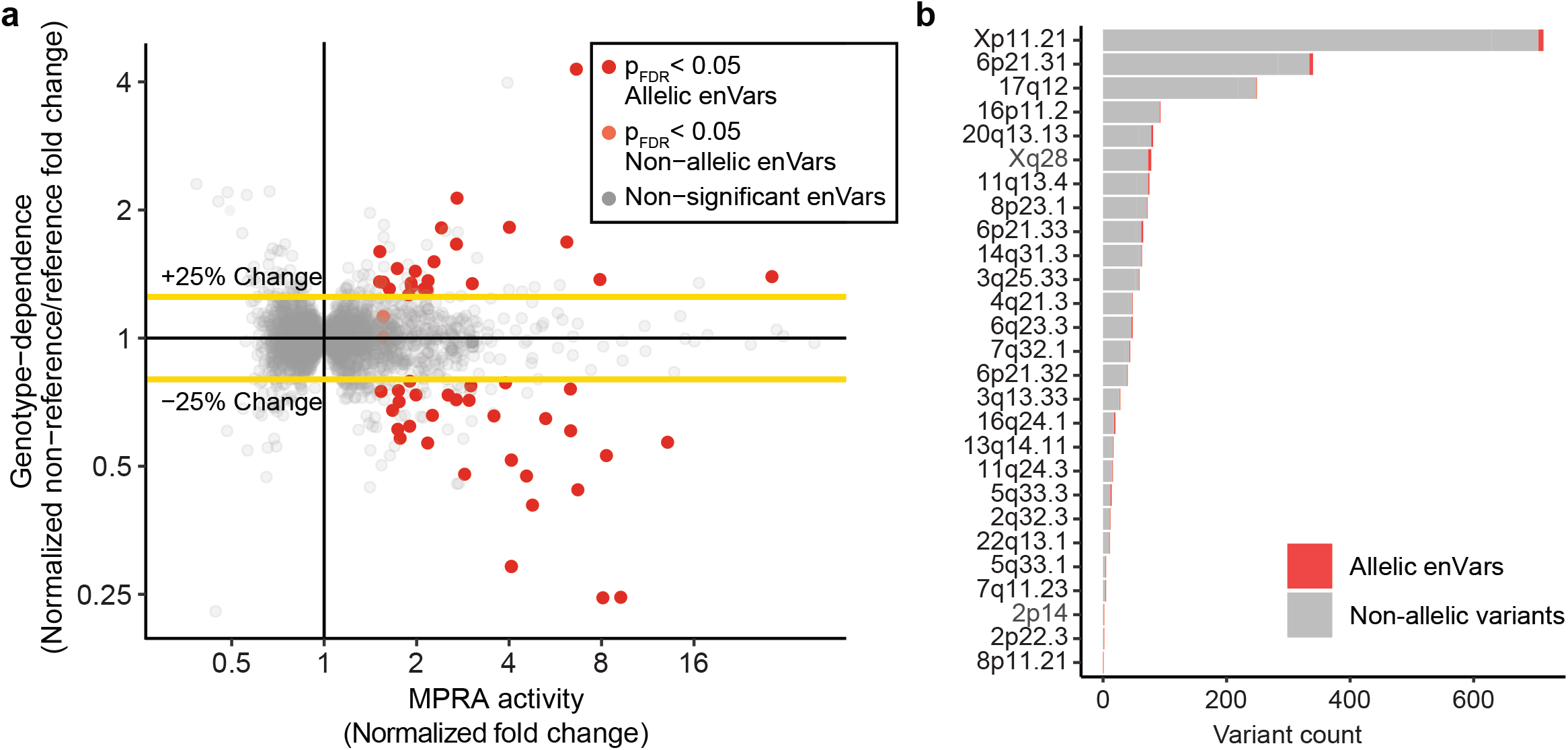
Regulatory activity of allelic enhancer variants (allelic enVars). a. Identification of allelic enVars. Genotype dependence (Y-axis) is defined as the normalized fold change of MPRA activity between the non-reference and reference alleles (see Methods). MPRA enhancer activity (X-axis) is presented as the maximum normalized fold-change of MPRA activity for any allele of the variant. Allelic enVars were defined as variants with a significant difference in MPRA activity (p_adj_ < 0.05) between any pair of alleles and at least a 25% change in activity difference (see Methods). b. MPRA enhancer activity at the 27 risk loci with at least one allelic enVar. Bar plots indicate the total number of variants at each locus. Variants with allelic enhancer activity (allelic enVars) are shown in red. Variants lacking allelic enhancer activity are shown in grey.

### Particular TFs have altered binding at SLE loci with allelic enhancer activity

To identify candidate regulatory proteins that might participate in allelic SLE mechanisms, we next used RELI to identify GM12878 ChIP-seq datasets that significantly overlap allelic enVars (**Supplemental Data Set 13**). Many of the top results are consistent with our previous study^10^, including the enriched presence of general enhancer features such as the H3K27ac histone mark (17 of 51 allelic enVars, 13.6-fold enriched, p_corrected_ <10^−38^), mediator complex subunit MED1 (17 of 51 allelic enVars, 13.0-fold enriched, p_corrected_ <10^−34^), and the histone acetyltransferase p300 (16 of 51 allelic enVars, 12.4-fold enriched, p_corrected_ <10^−32^), along with particular regulatory proteins that participate in “EBV super enhancers”^37^ and play key roles in B cells such as ATF7 (15 of 51 allelic enVars, 11.3-fold enriched, p_corrected_ <10^−26^), Ikaros/IKZF1 (19 of 51 allelic enVars, 9.7-fold enriched, p_corrected_ <10^−25^), and the NFκB subunit RELA (13 of 51 allelic enVars, 12.4-fold enriched, p_corrected_ <10^−24^). Also consistent with our previous study^10^, we observe strong enrichment for the EBV-encoded EBNA2 protein (7 of 51 allelic enVars, 17.7-fold enriched, p_corrected_ <10^−19^). Collectively, these data reveal particular regulatory proteins that might participate in the mechanisms contributing to SLE at multiple risk loci by driving allelic enhancer activity.

We next used the MARIO pipeline^10^ to search for allelic binding events (i.e. allelic imbalance between sequencing read counts) at SLE variants within 1,058 LCL ChIP-seq datasets (576 from GM12878). By necessity, this approach is limited to the 47 allelic enVars that are heterozygous in at least one of these cell lines. In total, this procedure identified 11 variants with strong allelic imbalance (MARIO ARS value > 0.4) in at least one ChIP-seq dataset (**Supplemental Data Set 14**), revealing groups of TFs and transcriptional regulators that allelically bind SLE risk variants with genotype-dependent MPRA enhancer activity. For example, the rs3101018 variant, which is associated with SLE^38^ and rheumatoid arthritis^39^ in Europeans, shows 1.7-fold stronger enhancer activity for the reference/non-risk ‘C’ allele compared to the non-reference/risk ‘T’ allele (**Figure 4A**). These results are consistent with a previously established eQTL obtained from GTEx^40^, which demonstrates higher Complement C4A (*C4A*) expression in EBV-transformed B cell lines for the rs3101018 ‘C’ allele than the ‘T’ allele (**Figure 4B**). Our MARIO allelic ChIP-seq analysis reveals 15 regulatory proteins that prefer the ‘C’ allele and 2 that prefer the ‘T’ allele (**Figure 4C, Supplemental Figure 4A**). Among these, particularly robust signal is obtained for ATF7, with one experimental replicate in GM12878 displaying 77 vs. 18 reads (‘C’ vs. ‘T’) and another showing 66 vs. 23 reads (‘C’ vs. ‘T’) (**Supplemental Data Set 14**). Moreover, CREB1 and CREM strongly favor the ‘C’ allele as well (**Supplemental Data Set 14**). In agreement with these data, computational analysis of the DNA sequences surrounding this variant predicts that ATF7, CREB1, and CREM will all bind more strongly to the ‘C’ than the ‘T’ allele (**Figure 4D**). Intriguingly, ten additional proteins (FOXK2, PKNOX1, ARID3A, ZBTB40, ZNF217, ARNT, ELF1, IKZF2, MEF2B, and FOXM1) also bind allelically and have known DNA binding motifs^41^, but none of them have binding sites altered by the variant. Further, we do not observe allelic chromatin accessibility in an available GM12878 ATAC-seq dataset (9 vs. 7 unique reads). Together, these results reveal a potentially causative SLE regulatory mechanism involving weaker direct binding of ATF7/CREB1/CREM to the ‘T’ risk allele, altering the recruitment of additional proteins to the locus and lowering the expression of *C4A*.

**Figure 4.**
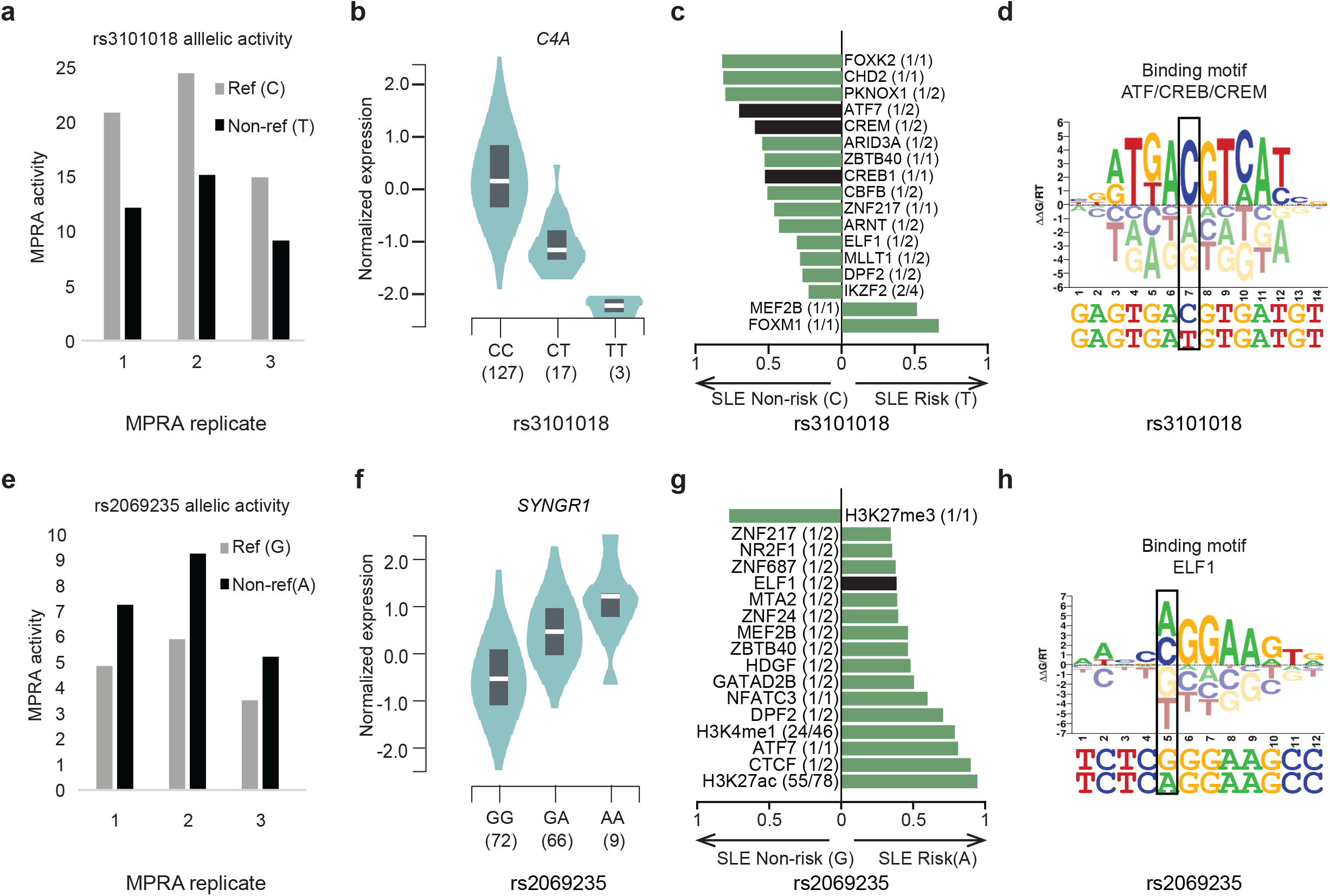
Lupus risk allele-dependent gene regulatory mechanisms at the *C4A* and *SYNGR1* genomic loci. a. and e. Normalized MPRA enhancer activity of each experimental replicate for rs3101018 and rs26069235. b. and f. Expression trait quantitative loci (eQTLs) revealing genotype-dependent expression of C4A and SNYGR1 for rs3101018 and rs26069235 in EBV-transformed B cell lines (GTEx). c. and g. Genotype-dependent activity of transcription factors, transcriptional regulators, and histone marks in EBV-transformed B cell lines for rs3101018 and rs26069235. Results with MARIO ARS value >0.4 and consistent allelic imbalance across ChIP-seq datasets are included (see Methods). The X-axis indicates the preferred allele, along with a value indicating the strength of the allelic behavior, calculated as one minus the ratio of the weak to strong read counts (e.g., 0.5 indicates the strong allele has twice the reads of the weak allele). The median value is plotted when data from multiple cell lines are available, with full results provided in **Supplemental Figure 4**. The numbers in parentheses represent the number of ChIP-seq datasets with significant allelic activity (i.e. MARIO ARS value > 0.4) out of the number of datasets where the given variant is inside a ChIP-seq peak and is also heterozygous in the given cell line. *Variant overlapping* TFs are indicated in black. *Variant adjacent* TFs are shown in green (see definition in **Figure 5a**). d. and h. DNA binding motif logos are shown for the ATF/CREB/CREM family, and ELF1 in the context of the DNA sequence surrounding rs3101018 and rs2069235, respectively. Tall nucleotides above the X-axis indicate preferred DNA bases. Bases below the X-axis are disfavored.

We observe a similar phenomenon for the rs2069235 variant, which is associated with SLE in Asian ancestries^42^ and rheumatoid arthritis in Europeans^43^. rs2069235 displays much stronger enhancer activity for the ‘A’ (non-reference/risk) allele compared to the ‘G’ (reference/non-risk) allele (**Figure 4E**), consistent with the established synaptogyrin 1 (*SYNGR1)* eQTL in EBV-transformed B cell lines^40^ (**Figure 4F**). Inspection of our allelic ChIP-seq data reveals 14 proteins preferentially binding the ‘A’ allele, with none preferring the ‘G’ allele (**Figure 4G, Supplemental Figure 4B**). Among these 14 proteins, only ELF1 has its binding site directly altered by the variant (**Figure 4H**). Strikingly, 55 of the 78 available H3K27ac datasets are allelic at this variant, with all 55 preferring the ‘A’ allele. Likewise, 24 of 46 H3K4me1 datasets are allelic, with all of them also preferring the ‘A’ allele. Both of these histone marks are indicative of active chromatin^28^. Only a single histone mark dataset prefers the ‘G’ allele – the H3K27me3 mark, which is indicative of silenced chromatin^28^ (**Figure 4G, Supplemental Figure 4B**). Together, these data are consistent with a potentially causative SLE molecular mechanism involving an allele-dependent enhancer consisting of stronger direct binding of ELF1 to the ‘A’ risk allele, along with indirectly altered binding of multiple additional TFs to this locus.

### Genotype-dependent binding to SLE variants with allelic enhancer activity by variant overlapping and variant adjacent TFs

As illustrated by the above examples, a particular TF can be involved in allelic mechanisms that are either directly impacted by a given variant (i.e. the variant directly alters the TF’s DNA-binding site) or indirectly impacted by the variant (i.e. the variant alters the DNA binding of the TF’s physical interaction partner, modulates chromatin accessibility, or affects another mechanism). At a given locus, we designate such TFs as *variant overlapping* and *variant adjacent* TFs, respectively (**Figure 5A**). We next sought to discover such TFs at the 51 allelic enVars. At each allelic enVar locus, we identified *variant overlapping* TFs as those TFs predicted to have strong binding to one allele and weak binding to the other allele. Likewise, we identified *variant adjacent* TFs as those TFs with proximal strong predicted binding sites that do not directly overlap the variant (see Methods). We then searched for particular TFs that tend to act as *variant overlapping* TFs or as *variant adjacent* TFs at the 51 allelic eVars using a proportion test (see Methods) and confirmed that their binding site locations are distributed relative to the variant as expected (**Figure 5B**). Consistent with our results at the *C4A* and *SYNGR1* loci, *variant overlapping* TFs include members of the ETS (e.g., ELF1) and ATF-like (e.g., ATF7) families, along with other TFs whose genetic loci are associated with SLE, including IRF5^44^ (**Figure 5C, Supplemental Data Set 15**). *Variant adjacent* TFs represent a largely distinct class, but also include several TFs with SLE genetic associations, including NFκB^45^, the Ikaros (IKZF) family^46^, and HMGA family members^47^ (**Figure 5D, Supplemental Data Set 15**). Collectively, these analyses reveal two distinct classes of TFs at a given SLE-associated locus that both likely play key roles in SLE mechanisms, along with particular TFs that tend to participate in one class or the other.

**Figure 5.**
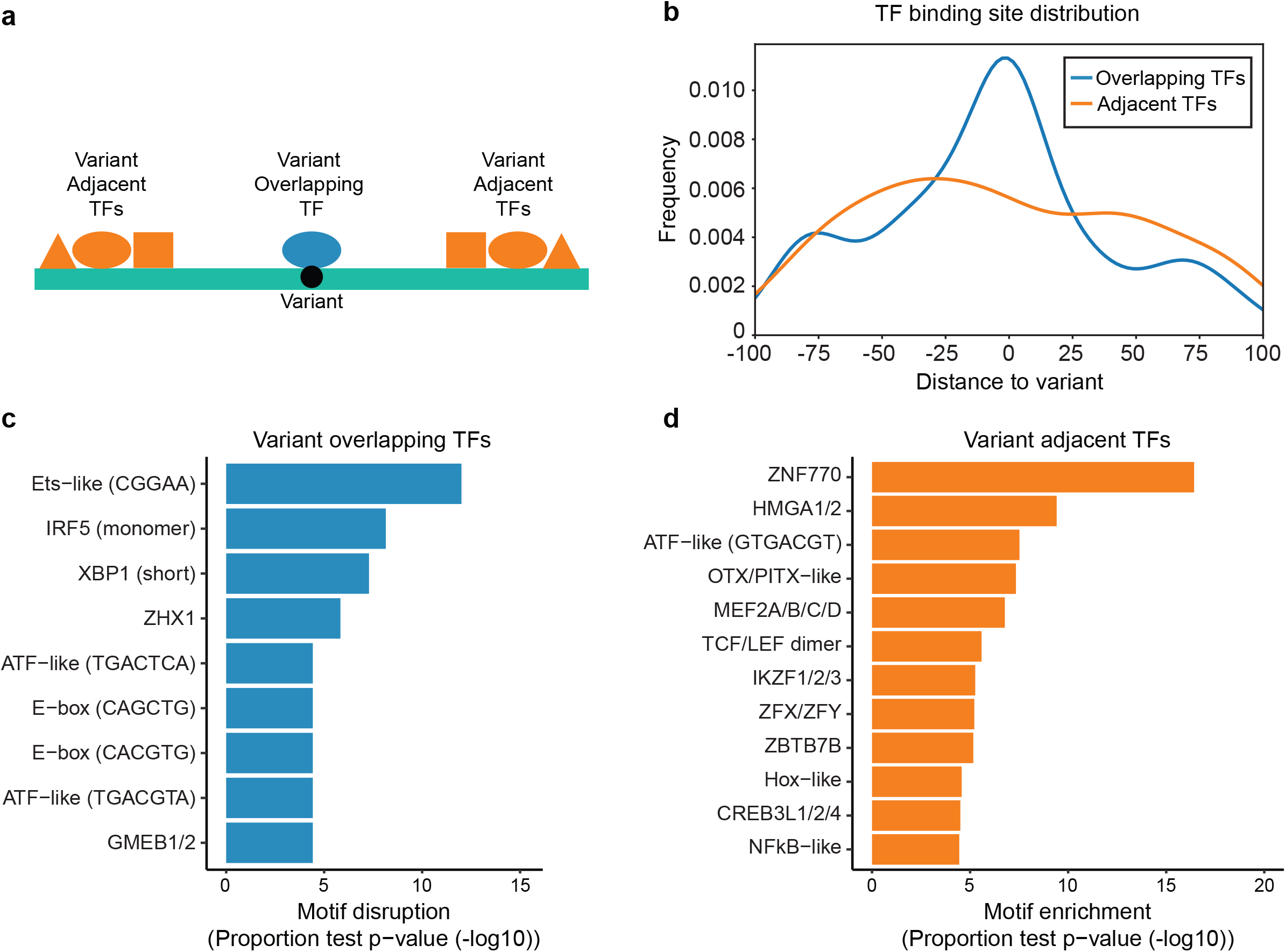
Identification of *variant overlapping* and *variant adjacent* TFs. a. Model of *variant overlapping* and *variant adjacent* transcription factors (TFs). *Variant overlapping* TFs allelically bind on top of variants, while *variant adjacent* TFs allelically bind near variants. b. TF binding site location distribution for *variant overlapping* and *variant adjacent* TFs, relative to allelic enVars. c. TF motif families enriched for participating as *variant overlapping* TFs at allelic enVars. Motif disruption p-values were estimated by comparing the fraction of motif disruption events at allelic enVars to the fraction observed at non-allelic enVars (see Methods). d. TF motif families enriched for participating as *variant adjacent* TFs at allelic enVars. Motif enrichment p-values were estimated by comparing the fraction of predicted TF binding sites in allelic enVars to random expectation (see Methods). For both the *variant overlapping* and *variant adjacent* analyses, motif families are shown with p_adj_ <0.0001 and three or more allelic events at allelic enVar loci, or five or more predicted binding sites at allelic enVar loci, respectively.

### Allelic transcription regulatory mechanisms shared and unique across cell types

To explore the cell-type specificity of allelic enhancer activity, we transfected our SLE MPRA library into Jurkat cells, a T cell line, as T cells are another important cell type in SLE^48^. We identified 92 SLE risk variants as allelic enVars in Jurkat cells, 25% of which were also found in GM12878 (**Supplemental Figure 5A, B, Supplemental Data Set 11**). We then repeated the experiment in Jurkat cells stimulated with the inflammatory cytokine TNFα, a key cytokine in SLE development^49^, to identify stimulation-dependent allelic enVars. This resulted in the identification of 102 allelic enVars, 28 of which were specific to the stimulated Jurkat cells (**Supplemental Figure 5C, D, Supplemental Data Set 11**). Altogether, our study identified a total of 145 allelic enVars across 50 independent SLE risk loci (**Supplemental Figure 5E**). These results highlight allelic transcriptional regulatory mechanisms that are both cell-type and inflammatory signaling dependent.

In summary, through the application of an allelic MPRA library to the EBV-transformed B cell line GM12878, we identified global transcriptional enhancer activity at 16% of SLE-associated genetic variants (enVars), with particular transcriptional regulatory proteins concentrated at these genomic locations. We further identified 51 SLE risk variants with allelic enhancer activity (allelic enVars) that we now nominate as plausibly causal by acting through genotype-dependent changes to enhancer activity in GM12878. Upon comparison to allelic enhancer activity in the Jurkat T cell line, we identified shared and unique allelic transcriptional regulatory mechanisms at SLE risk loci. Using experimental TF ChIP-seq data and TF binding site motif scanning, we propose a model where the collective action of the genotype-dependent binding of particular *variant overlapping* and *variant adjacent* TFs leads to genotype-dependent transcriptional activity at SLE risk loci.

## Discussion

Genome-wide association studies identify genetic loci with statistical disease associations. However, each risk locus often contains many plausibly causal variants due to linkage disequilibrium. This study is the first direct genome-wide measurement of enhancer activity at the ∼3,000 known SLE genetic risk variants in any context. Unbiased experimental approaches such as MPRA are vital for resolving causal variants and their molecular mechanisms of action.

Our results indicate that 16% of the SLE risk variants examined in this study have enhancer activity in the EBV-transformed B cell line GM12878. Furthermore, 51 of these enhancer variants at 27 loci have allelic enhancer activity. These findings are consistent with the theory that a large proportion of the genetic risk of SLE is mediated through transcriptional perturbation of critical B cell genes. Importantly, SLE risk loci exhibit cell-type and inflammatory signaling dependent allelic enhancer activity, with only 25% of allelic enVars shared between GM12878 and Jurkat cells. These results highlight both shared and unique allelic transcriptional regulatory mechanisms for SLE risk.

In this study, we used the EBV-transformed B cell line GM12878 as a model for exploring the effects of SLE risk variants. Previous studies have shown that B cells play a critical role in SLE development as immune cells that secrete autoantibodies driving etiology^11^. The relationship between EBV and SLE is widely appreciated. For example, EBV-infected B cells are more prevalent in SLE patients than in healthy people^13,14^ and patients with SLE have a higher EBV viral load and infection rate relative to controls^15,16^. Additionally, the EBV transcriptional regulator EBNA2 occupies SLE risk loci in a genotype-dependent manner^10^. *In vivo*, EBV infection can convert primary B cells to activated lymphoblasts^50,51^. EBV will eventually enter latency in resting memory B cells and establish lifelong infection^52^. *In vitro*, EBV infection transforms B cells into immortalized lymphoblastoid cell lines (LCLs)^17^. While we chose the EBV-transformed B cell line GM12878 as the primary disease model for our study, there are limitations to the use of EBV-transformed B cell lines. For example, GM12878 is an immortalized cell line with differences in DNA methylation and gene expression levels from resting and activated B cells^53,54^. A recent study also suggests that EBV infection causes B cells to undergo a germinal center-like differentiation into cells partially resembling plasmablasts and early plasma cells^55^, which is only a transient stage *in vivo*. Altogether, these limitations need to be considered when interpreting allelic transcriptional regulation of SLE risk variants in EBV-transformed B cell lines.

A critical finding of this study is that SLE risk variants with allelic enhancer activity likely alter the binding of many TFs. Although variants can directly affect the binding of *variant overlapping* TFs via disruption of a DNA-binding site, they can also simultaneously alter the binding of other *variant adjacent* TFs, presumably via genomic mechanisms such as altered chromatin accessibility, altered histone marks, indirect TF recruitment through physical interactions, changes in DNA shape, or changes to protein interaction partner DNA binding. This finding corroborates the previously proposed genetic variation-mediation model of motif-dependent and motif-independent TF binding^56-58^. In general, a given TF can be *variant overlapping* at one locus and *variant adjacent* at another, as exemplified by ATF7 (**Figure 4C, G**). Nonetheless, particular TF families tend to act as *variant overlapping* TFs at SLE loci (such as Ets, E-box, and ATF), whereas others tend to act as *variant adjacent* TFs (such as HMGA, Hox, and NFκB). Notably, many of these *variant overlapping* and *variant adjacent* TFs are themselves encoded by genetic risk loci associated with SLE (e.g., IRF5^44^, NFκB^45^, and ETS1^59^), suggesting that there are multiple means through which a particular TF can contribute to disease-based genetic mechanisms. For example, IRF5 targets might be mis-regulated in an SLE patient due to genetic associations in the promoter of *IRF5* that result in altered IRF5 protein levels^44^, or by genetic variants located within or adjacent to IRF5 binding sites at other genomic loci. It is currently unknown if these TF attributes are shared with other human diseases.

This study reveals possible causal genetic mechanisms involving altered binding of particular TFs at two important SLE risk loci. *C4A* is a component of the inflammatory complement pathway that is critical for the appropriate clearance of apoptotic cells^60^. People without *C4A* due to rare, protein-changing mutations are at a greatly increased risk for autoimmune diseases, including type I diabetes and SLE^61^. Further, the risk of developing SLE is 2.62 times higher in subjects with low total *C4A*^60^. Consistent with this observation, the SLE risk allele of rs3101018 at this genetic locus identified in our MPRA is associated with lower *C4A* expression (**Figure 4A**). Moreover, this variant is an eQTL for *C4A* in EBV-transformed B cell lines and whole blood cells, with the risk allele displaying lower *C4A* expression^62^ (**Figure 4B, Supplemental Data Set 7**). rs3101018 is located in the Human Leukocyte Antigen (HLA) region of the genome. While many genetic variants at this locus alter amino acid usage in major histocompatibility complex molecules and affect antigen presentation, non-coding genetic variants across the HLA region have also been demonstrated to affect gene expression independently of HLA-type^63-65^. Additionally, the SLE risk locus encoding *SYNGR1* was recently identified in a high-density genotyping study of subjects with Asian ancestry^42^ and also increases disease risk for schizophrenia^66^, primary biliary cirrhosis^67^, and rheumatoid arthritis^68^. *SYNGR1* is an integral membrane protein that is most robustly expressed in neurons of the central nervous system; however, there is measurable transcription and translation of *SYNGR1* in other tissues, including developing B cells^69^. eQTL data from EBV-transformed B cell lines and whole blood cells^62^ (**Figure 4F, Supplemental Data Set 7**) further support our MPRA-based findings of SLE risk genotype-dependent enhancer activity and gene expression at this locus. Altogether, the results of our MPRA study provide mechanistic insight into these variants through the identification of allelic enVars that facilitate SLE-risk genotype-dependent gene expression.

GWAS provides important discernment of the genetic origins of disease. In conjunction with other genome-scale assays such as ATAC-seq, ChIP-seq, and HiChIP-seq, MPRA reveals likely causal variants and genes, and enables the assembly of causal mechanisms affecting gene expression. In this study, we used MPRA to uncover specific genetic variants within the risk haplotypes of a complex disease in specific cell types. Our integrative analyses reveal specific molecular mechanisms underlying genotype-dependent transcriptional regulation and SLE disease risk. We conclude that MRPA is a robust tool for the nomination of causal genetic risk variants for any phenotype or disease with risk loci that act through genotype-dependent gene regulatory mechanisms, with this study providing a blueprint for dissecting the genetic etiology of many complex human diseases.

## Online Methods

### Variant selection and DNA sequence generation

All SLE-associated genetic risk loci reaching genome-wide significance published through March 2018 were included in this study^31,38,44,59,70-79^. A total of 91 genetic risk loci were used for linkage disequilibrium (LD) expansion (r^2^>0.8) based on 1000 Genomes Data^80^ in the ancestry(ies) of the initial genetic association using PLINK^81^ (**Supplemental Data Set 1**). All expanded variants were updated to the dbSNP 151 table^82^ from the UCSC table browser^83^ based on either variant name or genomic location. Unmappable variants were discarded. We also included 20 genetic variants from the Tewhey *et al*.^19^ study as positive and negative controls.

For single nucleotide polymorphisms, we pulled 170 base pairs (bps) of hg19-flanking DNA sequences for every allele, with the variant located in the center (84 bps upstream and 85 bps downstream of the variant). For the other types of variants (indels), we designed the flanking sequences to ensure that the longest allele has 170 bps. Adapters (15bps) were added to each sequence at either end (5’-ACTGGCCGCTTGACG − [170 bp oligo] - CACTGCGGCTCCTGC-3’) to make a 200 bp DNA sequence (**Supplemental Data Set 4**). For all resulting sequences, we created a forward and reverse complement sequence to compensate for possible DNA synthesis errors. A total of 12,478 oligos (3,093 variants, 6,239 alleles) were obtained from Twist Bioscience.

### MPRA experiments

#### Library Assembly

For assembly of the MPRA library, we followed the procedure described by Tewhey *et al*. ^19^ with minor modifications. In brief, we first created the empty vector pGL4.23∆xba∆luc from pGL4.23[luc2/minP] using primer Q5_deletion_rev and Q5_deletion_fwd following the manufacturer’s instruction of the Q5 Site-Directed Mutagenesis Kit. Then, 20bps barcodes were added to the synthesized oligos through 24X PCR with 50 μL system, each containing 1.86ng oligo, 25 μL NEBNext® Ultra™ II Q5® Master Mix, 1μM MPRA_v3_F and MPRA_v3_201_R. PCR was performed under the following conditions: 98°C for 2 mins, 12 cycles of (98°C for 10 sec, 60°C for 15 sec, 72°C for 45 sec), 72°C for 5 mins. Amplified product was purified and cloned into SfiI digested pGL4.23∆xba∆luc by Gibson assembly at 50°C for 1 hr. The assembled backbone library was purified and then transformed into *Escherichia coli* (*E*.*coli)* through electroporation (2kV, 200 ohm, 25 μF). Electroporated *E*.*coli* was expanded in 200 mL of LB Broth buffer supplemented with 100 μg/mL of carbenicillin at 37°C for 12 to 16 hrs. Plasmid was then extracted using the QIAGEN Plasmid Maxi Kit.

We next created the pGL4.23[eGFP/miniP] plasmid. An eGFP fragment was amplified from MS2-P65-HSF1_GFP (Addgene #61423) through PCR with a 50μl system containing 1 ng plasmid, 25 μL NEBNext® Ultra™ II Q5® Master Mix, 0.5 μM GFP_seq_MS2-P65-HSF1_GFP_FWD and GFP_seq_MS2-P65-HSF1_GFP_REV. PCR was performed under the following conditions: 98°C for 2 mins, 20 cycles of (98°C for 10 sec, 60°C for 15 sec, 72°C for 30 sec), 72°C for 5 mins. The amplified fragment was purified and then inserted into XbaI and NcoI digested pGL4.23[luc2/minP] through Gibson assembly at 50°C for 1 hr. The assembled plasmid was purified and then transformed into *E*.*coli* through chemical transformation. Transformed *E*.*coli* was expanded in 100 mL of LB Broth buffer supplemented with 100 μg/mL of carbenicillin at 37°C for 12 to 16 hrs. Plasmid was then extracted using the QIAGEN Plasmid Maxi Kit.

A miniP + eGFP fragment was amplified from pGL4.23[eGFP/miniP] through 8X PCR with 50 μL system, each containing 1 ng plasmid, 25 μL NEBNext® Ultra™ II Q5® Master Mix, 0.5μM 200-MPRA_v3_GFP_Fusion_v2_F and 201-MPRA_v3_GFP_Fusion_v2_R. PCR was performed under the following conditions: 98°C for 2 mins, 20 cycles of (98°C for 10 sec, 60°C for 15 sec, 72°C for 45 sec), 72°C for 5 mins. The amplified product was purified and then inserted into AsiSI digested backbone library through Gibson assembly at 50°C for 1.5 hrs to create the transfection library. The resulting library was re-digested by RecBCD and AsiSI, purified and then transformed into *E*.*coli* through electroporation (2kV, 200 ohm, 25 μF). Transformed *E*.*coli* was cultured in 5L of LB Broth buffer supplemented with 100 μg/mL of carbenicillin at 37 °C for 12 to 16 hrs. The plasmid was then extracted using the QIAGEN Endo-free Plasmid Giga Kit.

#### Oligo and barcode association

The oligo and barcode regions were amplified from the backbone library through 4X PCR with a 100 μL system containing 200 ng plasmid, 50 μL NEBNext® Ultra™ II Q5® Master Mix, 0.5 μM TruSeq_Universal_Adapter_P5 and MPRA_v3_TruSeq_Amp2Sa_F_P7. PCR was performed under the following conditions: 95°C for 20 sec, 6 cycles of (95°C for 20 sec, 62°C for 15 sec, 72°C for 30 sec), 72°C for 2 mins. The product was then purified, and indices were added through a 100μl system containing all purified product, 50μl NEBNext® Ultra™ II Q5® Master Mix, 0.5 μM TruSeq_Universal_Adapter_P5 and index primer. PCR was performed as above, except for only 5 cycles. Samples were purified, molar pooled, and sequenced using 2×125bp on Illumina NextSeq 500.

#### Transfection

The GM12878 cell line was grown in RPMI medium supplemented with 10% FBS, 100 units/mL of penicillin, and 100 µg/mL of streptomycin. Cells were seeded at a density of 5 × 10^5^ cells/mL the day before transfection. For triplicate transfections, we collected a total of 5 × 10^7^ cells per replicate. Cells were then suspended with 50 μg transfection library plasmid in 400 μL Buffer R. Electroporation was performed with the Neon transfection system in 100μl tips with 3 pulses of 1200V, 20 ms each. After transfection, cells were recovered in 50 mL pre-warmed RPMI medium supplemented only with 10% FBS for 24 hrs. Cells were then collected for preparation of the sequencing library for barcode counting.

The Jurkat cell line was grown in RPMI medium supplemented with 10% FBS, 100 units/mL of penicillin, and 100 µg/mL of streptomycin. Cells were seeded at a density of 5 × 10^5^ cells/mL the day before transfection. For each experimental group, we collected a total of 5 × 10^7^ cells per replicate for 5 replicates. Cells were then resuspended with 50 μg transfection library plasmid in 400 μL Buffer R. Electroporation was performed with the Neon transfection system in 100μl tips with 3 pulses of 1350V, 10 ms each. After transfection, cells were recovered in 50 mL pre-warmed RPMI medium supplemented only with 10% FBS for 24 hrs. After recovery, cells were supplemented with or without 100ng/ml TNF*α* for 24 hrs. Cells were then collected for preparation of the sequencing library for barcode counting.

#### Sequencing library for barcode counting

For sample from GM12878 cell, total RNA of transfected cells was extracted by the RNeasy Midi Kit following the manufacturer’s instruction. Extracted RNA was subjected to DNase treatment in a 375 μL system with 2.5 μL Turbo DNase and 37.5 μL Turbo DNase Buffer at 37°C for 1 hr. 3.75 μL 10% SDS and 37.5 μL 0.5M EDTA were added to stop DNase with 5 mins of incubation at 75°C. The whole volume was used for eGFP probe hybridization in an 1800 μL system, with 450 μl 20X SSC Buffer, 900 μL Formamide and 1 μL of each 100 μM Biotin-labeled GFP probe One to Three. The probe hybridization was performed through incubation at 65°C for 2.5 hrs. 200 μL Dynabeads™ MyOne™ Streptavidin C1 was prepared according to the manufacturer’s instruction. The beads were suspended in 250 μL 20X SSC Buffer and incubated with the above probe hybridization reaction at room temperature for 15 mins. Beads were then collected on a magnet and washed with 1X SSC Buffer once, and 0.1X SSC Buffer twice. eGFP mRNA was eluted first through adding 12.5 μL ddH2O, heating at 70°C for 2 mins and collecting on a magnet, then adding another 12.5 μL ddH2O, heating at 80°C for 2 mins and collecting on a magnet. All collected elution was performed with another DNase treatment in a 30 μL system containing 0.5 μL Turbo DNase and 3 μL Turbo DNase Buffer at 37°C for 1 hr. 0.5 μL 10% SDS was added to halt DNase reaction. Eluted mRNA was purified through RNA Clean SPRI Beads. mRNA was reverse transcribed to cDNA using SuperScript™ IV First-Strand Synthesis System with gene specific primer MPRA_v3_Amp2Sc_R, following the manufacturer’s instruction. cDNA and plasmid control were then used for building sequencing libraries following the Tag-seq Library Construction section in the paper of Tewhey *et al*.^19^. A total of two PCRs were needed for building the sequencing library. The first PCR was performed with TruSeq_Universal_Adapter_P5 and MPRA_V3_Illumina_GFP_F. The second PCR was performed with TruSeq_Universal_Adapter_P5 and index primer. Samples were purified, molar pooled, and sequenced using 1×75bp on Illumina NextSeq 500.

For samples from Jurkat cells, total DNA and RNA of transfected cells were extracted by the Qiagen ALLPrep DNA/RNA Mini Kit following the manufacturer’s instruction^23^. Extracted RNA was processed the same as above to obtain cDNA. cDNA, extracted DNA, and plasmid control were then used for building sequencing libraries following the Tag-seq Library Construction section in the paper of Tewhey *et al*.^19^. A total of two PCRs were needed for building the sequencing library. The first PCR was performed with TruSeq_Universal_Adapter_P5 and MPRA_V3_Illumina_GFP_F. The second PCR was performed with TruSeq_Universal_Adapter_P5 and index primer. Samples were purified, molar pooled, and sequenced using 1×100bp on Illumina NovaSeq 6000.

All primers used in this study are provided in **Supplemental Table 1**.

### MPRA data analysis

#### Oligo and barcode association

Paired-end, 125 bp reads were first quality filtered using Trimmomatic-0.38^84^ (flags: PE-phred33, LEADING:25, TRAILING:25, MINLEN:80). Read 1 was then separated into the 20bp barcode region and the oligo-matching region. The trimmed oligo-matching regions of Read 1 and Read 2 were mapped back to the synthesized oligo sequences using Bowtie2^85^ (flags: -X 250, ╌very-sensitive, -p 16). Barcodes were then associated with the oligo sequences using the read ID. Only uniquely mapped barcodes were used for downstream analysis.

#### Barcode counting

Single end 75/100 bp reads were first quality filtered using Trimmomatic-0.38^84^ (flags: PE - phred33, LEADING:3, TRAILING:3, MINLEN:70). Each read was then separated into the 20bp barcode region and the constant region. The trimmed constant regions of the reads were mapped back to the constant region within the eGFP 3’ UTR using Bowtie2^85^ (flags: ╌very-sensitive, -p 16). Only reads with Levenshtein distance of 4 or less within the constant region and perfect matches to the two bases directly adjacent to the barcode were kept. Barcodes were then associated with the retained reads using the read ID. Only barcodes that met our quality threshold requirements described above in the methods section “*Oligo and barcode association”* were used for downstream analysis.

#### Enhancer variant (EnVar) identification

We followed the procedures described in the “Identification of Regulatory Oligos” section of Tewhey *et al*.^19^ with minor modifications. In brief, oligos (alleles) with 30 or more unique barcodes from the plasmid control were included for analysis. All barcodes were summarized at the oligo level. Barcode count totals for each oligo, including all SLE variants and the 20 control variants, were passed into DESeq2^86^ (R version 3.5.3^87^) to estimate the fold change and significance between plasmid controls (see **Supplemental Note 1, Supplemental Figure 5F, G**) and the experiment replicates. A Benjamini-Hochberg FDR adjusted p-value of less than 0.05 was required for significance. Only significant alleles with greater than or equal to a 1.5x fold change were identified as enhancer alleles (enAlleles). A variant was identified as an enhancer variant (enVar) if any allele of this variant was an enAllele. Results for the 20 control variants were compared to data from Tewhey *et al*.^19^ to estimate accuracy, sensitivity, and specificity.

#### Allelic enVar identification

Only enVars were considered for allelic analysis. The barcode counts from every allele of each enVar were used for calculating p-values by comparing the log2 ratios of the non-reference allele vs the reference allele, normalized by plasmid controls, using Student’s *t*-test^19,35^. p-values were adjusted with the Benjamini-Hochberg FDR-based procedure. A corrected p-value of less than 0.05 was required for significance. Only significant alleles with 25%-fold changes or greater were identified as allelic enVars (see **Supplemental Note 2, Supplemental Figure 6**). We have created an R package (mpraprofiler) for performing this analysis, which is available on the Weirauch lab GitHub page (https://github.com/WeirauchLab/).

### Downstream MPRA analysis

#### Gene annotation

We annotated each SLE genetic variant with its nearest gene using the NCBI RefSeq table^88^ downloaded from the UCSC table browser^83^. enVars were annotated using a combination of DNA looping interactions (GM12878 Capture Hi-C data^89,90^) and eQTL data obtained from the eQTL Catalogue, a resource that contains quality controlled, uniformly re-computed eQTLs from 19 eQTL publications^62,91-110^, EBV-transformed B cell lines (GTEx Analysis V7 (dbGaP Accession phs000424.v7.p2)^40^ and other individual studies^111-114^. For all variants, the target genes were annotated (**Supplemental Data Set 2**) using the union of promoter interacting genes and eQTL genes from B cells with and without EBV transformation, when available. Otherwise, target genes were annotated as the nearest gene. Allelic enVar gene targets were classified into four tiers: a Tier 1) variant is both an eQTL and also loops to the promoter of the same gene; a Tier 2) variant has an eQTL for at least one gene; a Tier 3) variant only loops to the promoter of at least one gene; a Tier 4) variant is neither an eQTL nor loops to the promoter of any gene (**Supplemental Data Set 12**).

#### TF binding site motif enrichment analysis

To identify specific TFs whose binding might contribute to the enhancer activity observed in our MPRA experiments, we performed HOMER^33^ TF binding site motif enrichment analysis. Specifically, we used HOMER to calculate the enrichment of each motif in the sequence of enAlleles compared to the sequences of non-enAlleles. HOMER was modified to use the large library of human position weight matrix (PWM) binding site models contained in build 2.0 of the CisBP database^34^ and a log base 2 likelihood scoring system.

#### GO enrichment analysis

Enrichr^115,116^ was used for GO enrichment analysis. In short, the target genes of enVars were passed to Enrichr for analysis. Results from the GO biological process (2018) category were used (**Supplemental Data Set 8, Supplemental Figure 3)**.

#### Identification and processing of publicly available LCL ChIP-seq data

1,058 ChIP-seq datasets were obtained from the Gene Expression Omnibus (GEO)^117^ using custom scripts that searched for ChIP-seq experiments performed in EBV-transformed lymphoblastoid cell lines (LCLs). The annotations for every dataset (assay type, cell line, assayed molecule) were manually checked by two authors (MTW and LCK) to ensure accuracy. The Sequence Read Archive (SRA) files obtained from GEO were analyzed using an automated pipeline. Briefly, the pipeline first runs QC on the FastQ files containing the sequencing reads using FastQC (v0.11.2)^118^. If FastQC detects adapter sequences, the pipeline runs the FastQ files through Trim Galore (v0.4.2)^119^, a wrapper script that runs cutadapt (v1.9.1)^120^ to remove the detected adapter sequence from the reads. The quality controlled reads are then aligned to the reference human genome (hg19/GRCh37) using bowtie2 (v2.3.4.1)^85^. The aligned reads (in .BAM format) are then sorted using samtools (v1.8.0)^121^ and duplicate reads are removed using picard (v1.89)^122^. Finally, peaks are called using MACS2 (v2.1.2) (flags: callpeak -g hs -q 0.01 -f BAM)^123^.

ENCODE blacklist regions^124^ were removed from the peak sets using the hg19-blacklist.bed.gz file available at https://github.com/Boyle-Lab/Blacklist/tree/master/lists/Blacklist_v1. ChIP-seq datasets GSM1666207, GSM2748907 and GSM1599157 were removed due to the low number of cells used in the experiments.

#### Functional genomics dataset enrichment analysis with RELI

We used the RELI^10^ algorithm to identify genomic features (TF binding events, histone marks, etc.) that coincide with enVars. As input, RELI takes the genomic coordinates of enVars. RELI then systematically intersects these coordinates with one of the GM12878 ChIP-seq datasets, and the number of input regions overlapping the peaks of this dataset (by at least one base) is counted. Next, a p-value describing the significance of this overlap is estimated using a simulation-based procedure. To this end, a ‘negative set’ is created for comparison to the input set, which in this study contains the set of non-enVars (i.e., variants with no allele having an adjusted p-value of less than 0.05 and more than 10%-fold change in the DESeq2 result). A distribution of expected overlap values is then created from 2,000 iterations of randomly sampling from the negative set, each time choosing a set of negative examples that match the input set in terms of the total number of genomic loci. The distribution of the expected overlap values from the randomized data resembles a normal distribution and can thus be used to generate a Z-score and corresponding p-value estimating the significance of the observed number of input regions that overlap each ChIP-seq dataset.

We performed similar RELI analysis for allelic enVars. As input, we used the allelic enVar sites. For the ‘negative set’, we used the set of common SNPs taken from the dbSNP142 database downloaded from the UCSC table browser^83^.

#### Identification of allelic ChIP-seq reads using MARIO

To identify possible mechanisms underlying our allelic enVars, we applied our MARIO^10^ method to the LCL ChIP-seq dataset collection described above. In brief, MARIO identifies common genetic variants that are (1) heterozygous in the assayed cell line and (2) located within a peak in a given ChIP-seq dataset. It then examines the sequencing reads that map to each heterozygote in each peak for imbalance between the two alleles. Results are combined across experimental replicates to produce a robust Allelic Reproducibility Score (ARS). Results with MARIO ARS value > 0.4 that also pass the following three post-processing filters were considered allelic. (1) The variant must be significantly allelic for a given protein/histone mark (ARS > 0.4) in at least 50% of the datasets in which that variant was heterozygous; (2) The same allele must be significantly preferred (ARS > 0.4) in at least 75% of the datasets where that variant shows significant allelic behavior; and (3) The replicates of a given experiment must all prefer the same strong allele. These post-processing filters were applied to remove results with inconsistent allelic imbalance, extending the procedures of our previous study^10^.

#### Identification of variant overlapping and variant adjacent TFs

*Variant overlapping* TFs were identified using an algorithm that compares predicted TF binding motif scores between the different alleles of each allelic enVar. First, we padded each allele of a given allelic enVar with 25 bps of upstream and downstream DNA sequence (a sufficient length to account for any known human TF binding sites^125^). The algorithm consists of two major components: (1) individually scoring the two alleles of a given variant with a given TF model; and (2) quantifying the difference in the binding intensity between these two alleles. DNA sequences are scored using the large collection of human TF position frequency matrix (PFM) models contained in the Cis-BP database^34^ and the log-likelihood PFM scoring system^126^. Since log-likelihood score distributions vary substantially (depending on the information content of a given motif), we employ a simple scaled scoring system that maps a given log-likelihood score to the percentage of the maximum achievable log-likelihood score of the given motif – we refer to this value as the “relative PFM score”. We identify binding site altering events (i.e. “creating” or “breaking” a predicted binding site for a given TF motif) as cases where one allele has a relative PFM score of 70% or higher, and the other allele has a score of less than 40%. For a given variant, any TF with allelic ChIP-seq sequencing reads (see above) and a binding site altering event for any of its motifs was deemed a *variant overlapping* TF. Any TF with allelic ChIP-seq sequencing reads and a lack of a binding site altering event for any of its motifs was deemed a *variant adjacent* TF.

We next sought to identify particular TFs that tend to be *variant overlapping* TFs at SLE allelic enVars. To this end, we calculated the fraction of times each TF motif has a binding site altering event (as defined above) at SLE allelic enVars. As background, we calculated the fraction of times each TF motif has a binding site altering event at non-allelic enVars. The significance of the difference between these two fractions was then calculated using a proportions test. Results are provided in **Supplemental Data Set 15**.

We used a similar procedure to identify particular TFs that tend to be *variant adjacent* TFs at SLE allelic enVars. We performed HOMER motif enrichment analysis using the full 170bp allelic enVar DNA sequences as input. The standard HOMER null model (scrambled input sequence, maintaining dinucleotide frequencies) was used as background. The fractions of motif “hits” obtained in the foreground vs. background set were then compared, and significance was again calculated using a proportions test. Results are provided in **Supplemental Data Set 15**.

## Data availability

MPRA data are available in the Gene Expression Omnibus (GEO) database under accession number GSE143792. Full datasets and processed results are provided in the Supplementary Material.

## Code availability

Source code, with full documentation and examples, are freely available under the GNU General Public License on the Weirauch Lab GitHub page: https://github.com/WeirauchLab

## Acknowledgements

We thank Roger Pique-Regi and Francesca Luca for their consultation and guidance in the development of our MPRA libraries and approach. We thank Kevin Ernst for computational support. We greatly appreciate Aaron Zorn, Raphael Kopan, Marc Rothenberg, and Stephen Waggoner for constructive feedback and guidance.

This work was funded by the National Institutes of Health (F32 AI129249, K99-HG009920, P30 AR070549, P30 DK078392, R01 AI024717, R01 AI148276, R01 AR073228, R01 DK107502, R01 GM055479, R01 HG010730, R01 NS099068, U01 AI130830, U01 HG008666), Cincinnati Children’s Hospital Research Foundation (Academic and Research Committee award, Center for Pediatric Genomics pilot funding, Endowed Scholar award), the Veterans Administration (I01 BX001834), the Ohio Supercomputing Center, and the State of Ohio.

## Author Contributions

This manuscript was written by XL, XC, MTW, and LCK with critical input from CF, OD, DM, SP, TH, YH, MP, TC, ERM, JPR, CGdB, and JBH. Experiments were designed by XL, JPR, CGdB, MTW, and LCK with computational design by XL, XC, SP, JPR, CGdB, MTW, and LCK. Experiments were performed by XL, CF, OD, DM, and YH. Analysis was performed by XL, XC, and SP. Data were provided by TH, MP, TC, ERM, and MTW. Funding was provided by JPR, CGdB, JBH, MTW, and LCK.

