## Supplemental Note for "Genome-wide discovery of SLE genetic risk variant allelic enhancer activity"

**Supplemental Note 1. Normalization control optimization**

To our knowledge, two methods have been used to date as normalization controls for mRNA barcode counts: plasmids used for transfection[1] and DNA extracted from transfected cells[2]. We compared the allelic enVars that we identified through these two normalization methods using MPRA data from Jurkat cells with and without TNF$\alpha$ simulation. All MPRA data analysis steps were identical in this comparison, other than the use of plasmid or extracted DNA as the normalization control. As shown in **Supplemental Figure 5F, G,** there is little difference in the results produced by the two different normalization strategies. Moreover, the plasmid control strategy identified slightly more allelic enVars in both experiments. We conclude from these comparisons that the plasmid control is an effective, and slightly more sensitive normalization approach.

**Supplemental Note 2. Comparison of MPRA data analysis methods**

To date, four methods have been used to our knowledge for enAllele/enVar and allelic enVar identification: DESeq2 combined with Student's *t*-test[1,3], QuASAR-MPRA[4], mpralm[5] and MPRAnalyze[6]. For the DESeq2 combined with Student's *t*-test method, we followed the analysis procedure in Tewhey *et al.*[1], with the application of additional more stringent criteria, as described in the "*Enhancer variant (EnVar) identification*" and *"Allelic enVar identification"* sections of the Methods. For the other three methods, we followed the analysis procedures described in the corresponding publications.

To compare the performance of these four methods, we developed approaches to compare their (1) specificity, (2) consistency, and (3) sensitivity. We note that this analysis was hindered somewhat due to the lack of a reliable gold standard. First, we combined all of our GM12878 experimental replicate data and plasmid control data and re-sampled them into four replicate sets (repA, repB, repC, and repD), respectively. We then grouped repA and repB into Group 1 and repC and repD into Group 2. Next, we ran each of the four analysis methods to identify enAlleles and allelic enVars in Group 1 and Group 2. We expected to find a small number of differences between the results of Group 1 and Group 2, since they are sampled from the same set of reads.

Of the four analysis methods, DESeq2 combined with Student's *t*-test, QuASAR-MPRA and mpralm methods ran successfully in our hands. The MPRAnalyze pipeline failed to run and had extensive computational resource requirements - the program ran with 200GB of RAM for >17hrs, with no successful completion in three attempts.

To assess specificity, we compared the Group 1 and Group 2 enAllele predictions. Both the DESeq2 combined with Student's *t*-test and mpralm methods found no significant differences between the subsampled groups, indicating strong specificity (and also further highlighting the reproducibility of our data). QuASAR-MPRA does not have a function to perform enhancer identification, so we could not gauge its specificity using this procedure.

To assess consistency, we identified allelic enVars in Group 1 and Group 2, respectively and used a $p<0.05$ allelic cut-off. From a total of 3,093 variants, the DESeq2 combined with Student's *t*-test method identified a total of 83 allelic enVars in the two groups, with 57% in common between Group 1 and Group 2. QuASAR-MPRA identified 39 allelic enVars, with 92% in common between Group 1 and Group 2. mpralm identified 1,114 allelic enVars, with 49% in common between Group 1 and Group 2 (**Supplemental Figure 6A**). 1,114 allelic enVars is 36% of all tested variants, which indicated to us that a more stringent p-value cut-off was needed for mpralm. The number of allelic enVars dropped exponentially at more stringent cut-offs (**Supplemental Figure 6B**). Based on this analysis, we chose a cut-off $p<10^{-10}$, which results in 94 allelic enVars in the two groups (a comparable number to that identified by the other methods). At this threshold, only 17% of the mpralm-identified allelic enVars were shared in both groups (**Supplemental Figure 6C**). Additionally, because Group 1 and Group 2 were sampled from the same pool, it was concerning that mpralm consistently identified more significant allelic enVars from Group 1 compared to Group 2 despite re-running of the code numerous times by two different people. This inconsistency combined with the inflated number of identified allelic enVars at reasonable p-value cutoffs greatly reduced our enthusiasm for mpralm.

Considering the poor performance of mpralm, we subsequently focused on the DESeq2 combined with Student's *t*-test and QuASAR-MPRA methods. We first compared the final allelic enVar predictions of these two methods (i.e. those predicted by the given method to be allelic in both Group 1 and Group 2). As shown in **Supplemental Figure 6D**, DESeq2 combined with Student's *t*-test identified 27 unique variants, while QuASAR-MPRA identified 16 unique variants. The unique QuASAR-MPRA variants tended to have a larger variance between replicates, diminishing our confidence in these results. Variants unique to the DESeq2 combined with Student's *t*-test method tend to have a much wider range of genotype-dependence values (**Supplemental Figure 6E**), indicating that this method is likely more sensitive. To further explore sensitivity, we manually examined raw MPRA data for four unique Student's *t*-test variants (the two variants with the largest amount of genotype-dependence and the two with the smallest). As shown in **Supplemental Figure 6F**, all four variants show consistent allelic difference among the replicates in both groups, raising questions about why QuASAR-MPRA did not detect them. Taken together, these results suggest to us that the DESeq2 combined with Student's *t*-test method currently represents the analytical strategy with the best balance between specificity, consistency, and sensitivity.
